# Deep learning microstructure estimation of developing brains from diffusion MRI: a newborn and fetal study

**DOI:** 10.1101/2023.07.01.547351

**Authors:** Hamza Kebiri, Ali Gholipour, Lana Vasung, Željka Krsnik, Davood Karimi, Meritxell Bach Cuadra

## Abstract

Diffusion-weighted magnetic resonance imaging (dMRI) is widely used to assess the brain white matter. Fiber orientation distribution functions (FODs) are a common way of representing the orientation and density of white matter fibers. However, with standard FOD computation methods, accurate estimation of FODs requires a large number of measurements that usually cannot be acquired for newborns and fetuses. We propose to overcome this limitation by using a deep learning method to map as few as six diffusion-weighted measurements to the target FOD. To train the model, we use the FODs computed using multi-shell high angular resolution measurements as target. Extensive quantitative evaluations show that the new deep learning method, using significantly fewer measurements, achieves comparable or superior results to standard methods such as Constrained Spherical Deconvolution. We demonstrate the generalizability of the new deep learning method across scanners, acquisition protocols, and anatomy on two clinical datasets of newborns and fetuses. Additionally, we compute agreement metrics within the HARDI newborn dataset, and validate fetal FODs with post-mortem histological data. The results of this study show the advantage of deep learning in inferring the microstructure of the developing brain from in-vivo dMRI measurements that are often very limited due to subject motion and limited acquisition times, but also highlight the intrinsic limitations of dMRI in the analysis of the developing brain microstructure. These findings, therefore, advocate for the need for improved methods that are tailored to studying the early development of human brain.

## Introduction

Early brain growth is characterized by rapid and complex structural and functional developments that are vulnerable to various genetic and environmental factors. The influence of early brain development and disorders on the brain health later in life has received growing interest^1–5^. Magnetic Resonance Imaging (MRI) is a non-invasive method for assessing these developments *in vivo*. Diffusion MRI (dMRI), specifically, offers a means to assess the micro-structure of the white matter using the diffusion of water molecules as a proxy measure^6,7^. However, application of dMRI to study the developing brain has been limited due to motion, limited scan time, and low signal-to-noise ratio (SNR)^8–10^. Despite these limitations, prior works have shown the potential of dMRI to probe the early brain development. For instance, several studies^11–13^ have used spatiotemporal changes in Fractional Anisotropy (FA), Mean Diffusivity (MD) and different cortical morphology indices to characterize the normal brain development. Recent availability of large high-quality datasets such as those collected under the developing Human Connectome Project (dHCP)^14,15^ present a unique opportunity to enhance our understanding of the developing brain. These datasets include dense multi-shell measurements. As such, derived dMRI quantities can be considered as reference values or gold standards to which derived metrics from more constrained clinical datasets, which usually do not exceed 15 diffusion measurements with a single low b-value (500 − 750*s/mm*^2^), can be compared.

The prevailing way of extracting diffusion properties from the diffusion signal involves a model, typically a diffusion tensor imaging (DTI) model^16^. More complex models such as the multi-shell multi-tissue constrained spherical deconvolution (MSMT-CSD)^17,18^ aim to reconstruct Fiber Orientation Distribution Functions (FODs) that allow depiction of more intricate white matter configurations such as fiber crossings. These models require densely sampled multi-shell data. The output of all of these models can be studied directly, i.e. by computing metrics such as FA or MD from the diffusion tensor or the apparent fiber density^19^ from the FOD. Alternatively, they can be further processed globally to reconstruct the fiber tracts^20,21^ that are responsible for transmitting action potentials between different regions of the brain.

In general, mapping the acquired diffusion signal to an interpretable and informative diffusion metric requires a prior model. Conventional estimation methods do not provide feedback regarding which measurements are more informative for estimating the given model. Differently, deep neural networks can treat the problem as a single learnable task that can be optimized via *back-propagation*, by directly learning a mapping between the diffusion signal and the target diffusion quantity. Hence, bypassing the sub-optimal model fitting step that can be sensitive to noise. Golkov et al.^22^ proposed the first deep learning (DL) model that directly estimated diffusion kurtosis^23^ and neurite orientation dispersion and density measures^24^ from a small number of diffusion measurements in adult brains. They showed a drastic decrease in scanning time with limited loss in accuracy. Since then several other works have explored DL methods in adult brains as to directly estimate diffusion scalars or model reconstruction (FODs). For instance, with superDTI^25^, accurate predictions of tensor maps using a neural network were achieved using only six diffusion measurements. This model was robust to various noise levels and could depict lesions present in the dataset. Kopper et al.^26^, employed a 2D convolutional neural network (CNN) in a classification approach to predict the orientation of fibers, while Lin et al.^27^ utilized a 3D CNN to predict FODs based on a small neighborhood of the diffusion signal. Karimi et al.^28^ used a multi-layer perceptron to predict FODs. To leverage correlations between neighboring voxels,^29^ used a two-stage Transformer-CNN to map 200 measurements to 60 measurements, followed by predicting FODs. Nonetheless acquiring such a large number of measurements is difficult and frequently infeasible for noncooperative cohorts, such as neonates or fetuses.

The challenge of acquiring useful dMRI data from newborns and fetuses who tend to unpredictably move during long and loud dMRI acquisitions is exacerbated by the low signal available from the small size of the immature and developing brain compartments, the low resolution of dMRI, and the rapid and large changes that occur to the brain microstructure across gestation and early after birth.

These complex maturation processes that are unfolding during gestation include the development of major fiber bundles, namely limbic and projection fibers during the first trimester^30^. For instance, the internal capsule experiences intricate microstructure alterations as a result of the intertwining of multiple fiber pathways that initiate development during different periods of gestation. In the second trimester, association fibers start developing and become evident at the third trimester. Specifically, the superior longitudinal fasciculus exhibits accelerated growth during this phase and continues to undergo substantial development even beyond the time of birth^31^. The radial coherence within the telencephalic wall gradually diminishes with gestational weeks (GW). Furthermore, the regional radial coherence within the deep subplate zone starts to vanish around 26 GW. This regional loss of radial coherence aligns temporally with the previously reported emergence of long association cortico-cortical tracts^32^.

These dynamic changes and the other aforementioned problems of this sensitive population, added to B0 and B1 inhomogeneities, pose additional challenges for learning-based methods for FOD estimation such as those reviewed above. Therefore, in this study we aimed to investigate the use and generalizability of deep learning to estimate the microstructure of the developing brain in newborns and fetuses.

To the best of our knowledge, these learning-based FOD estimation methods have not yet been critically evaluated for fetal populations and in non-research protocols of newborns. In this study, we demonstrate that a deep convolutional neural network with a large field of view (FOV) can accurately estimate FODs using only 6-12 diffusion-weighted measurements. Firstly, we show, on N=465 subjects from the dHCP dataset, that a deep learning approach can achieve a level of accuracy that is comparable to the accuracy of the state-of-the-art methods, while reducing the required number of measurements by a factor of ∼21-43. Secondly, we present evidence of a low agreement among standard FOD estimation methods for these age groups. Thirdly, we show the generalizability of deep learning methods on two out-of-domain clinical datasets of 26 *in vivo* fetuses and neonates that were scanned with different scanners and acquisition protocols. Finally, we assess for the first time, the deep learning generated fetal FODs with post-mortem histological data of corresponding gestational weeks.

## Results

### Research dMRI acquisitions of neonates

We trained our deep learning model (*DL*_*n*_) on dMRI data from neonates. FODs estimated with the MSMT-CSD method using 280 diffusion-weighted and 20 b0 measurements were used as estimation target. We refer to MSMT-CSD estimations as ground truth (GT). The input to *DL*_*n*_ consists of 6 diffusion-weighted measurements normalized with one *b0* measurement. After training, the network was applied on independent test data.

Qualitatively, the FODs estimated by *DL*_*n*_ were very similar to those estimated by MSMT-CSD using 300 measurements in 3 shells (Figure 1). We also compared our results with those of CSD using 148 measurements. CSD overestimates the number of peaks in the regions outside of deep white matter. Although estimating the 45 FOD coefficients using 6 measurements is an under-determined problem, for the sake of comparison we present the CSD estimated FODs with the six measurements as those used for *DL*_*n*_ (CSD-6 in Figure 1). CSD-6 results show significant errors even in the location of major white matter tracts such as the corpus callosum.

**Figure 1.**
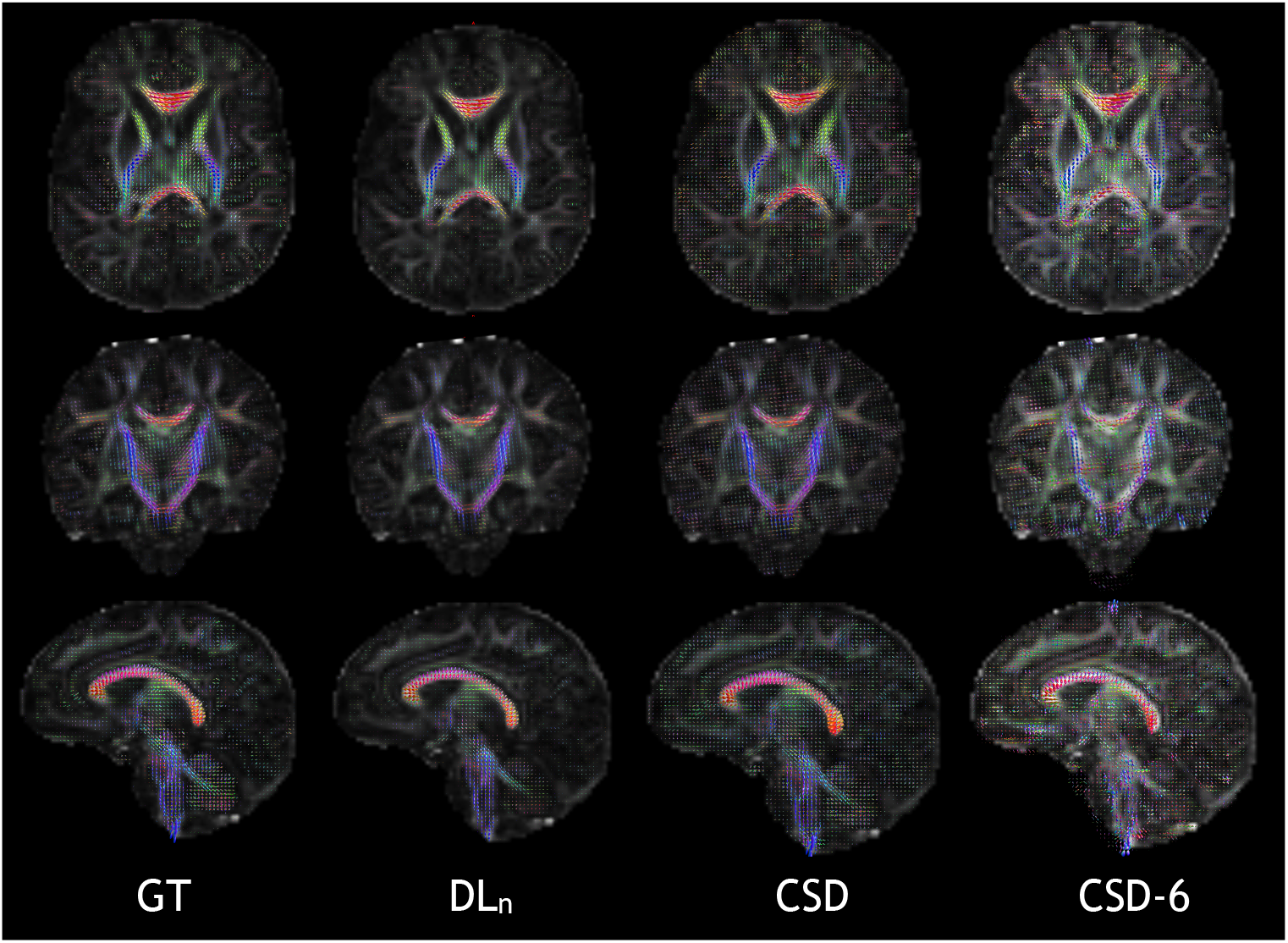
Qualitative high level comparison between, from left to right, the MSMT-CSD GT using the 300 multi-shell samples, the deep learning method *DL*_*n*_ using six b = 1000 *s/mm*^2^ measurements and one *b0*, and CSD using 128 measurements of b = 2600 *s/mm*^2^ and 20 *b0* images. Also shown on the right CSD-6, i.e. CSD with the same measurements that the DL method used. Axial, coronal and sagittal views are shown from top to bottom and the background images correspond to fractional anisotropy (FA) extracted from diffusion tensors estimated with all b = 1000 *s/mm*^2^ measurements.

#### Quantitative assessment

On N=320 independent test subjects from the dHCP dataset, *DL*_*n*_ showed low estimation error (with respect to the MSMT-CSD GT) in terms of several metrics compared to the various standard estimation methods. We assessed the reproducibility of the GT by applying MSMT-CSD on subsets of the measurements from the same subject. Specifically, we split the 300 measurements into two disjoint subsets of 150 multi-shell measurements and applied MSMT-CSD on each subsets to compute two independent FOD estimations, which we denote with GS1 and GS2. GS1 and GS2 can be viewed as two high-quality scans of the same subject, conducted with a similar protocol.

As depicted in Figure 2, the *DL*_*n*_ model has lower error rates on apparent fiber density (AFD)^19^ of 0.178 (±0.083) with respect to the GT, which is close to the corresponding gold standard difference of 0.064 (±0.034), compared to all other techniques. The other methods, namely Constrained Spherical Deconvolution (CSD)^17^ using 148 measurements, Constant Solid Angle^33^ (CSA), and the Sparse Fascicle Model (SFM) using all 300 measurements, displayed elevated error rates (both in terms of means and standard deviations) than the *DL*_*n*_ model. It is noteworthy that statistically significant differences were observed between the *DL*_*n*_ and the other methods with *p* ≤4.8^−11^ based on paired t-test with Bonferroni correction for multiple comparisons.

**Figure 2.**
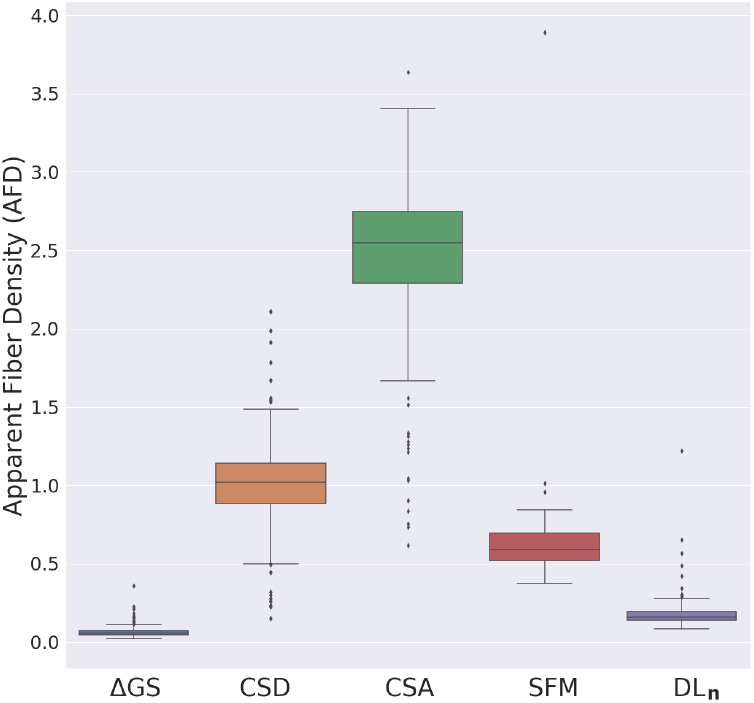
Apparent fiber density error with respect to the MSMT-GT for the different methods, along with the agreement between the two gold standard datasets (Δ*GS*) that is shown as an upper bound error. The different baseline methods used are Constrained Spherical Deconvolution (CSD)^17^, using 128 gradient directions (b-value of 2600 *s/mm*^2^) and 20 *b0* images; Constant Solid Angle^33^ (CSA) and the Sparse Fascicle Model (SFM) model^34^ using all available 300 measurements. *DL*_*n*_ method, with less than an order of magnitude in the number of samples (six b=2600 *s/mm*^2^ samples) and one *b0* image) achieves the lowest error by a high margin.

Peaks count and orientations were also estimated from the FOD generated by all methods. We first identified a low level of agreement rate (AR) for multiple fibers within the MSMT-CSD GT (Δ*GS*) as can be shown in Figure 3 (a). The AR was extracted from the confusion matrix of the estimated number of peaks (Table 1, details in Methods section). For instance, the 1-peaks AR was 88.2%, while 46.7% and 47.2% were observed for 2-peaks and 3-peaks, respectively. Our proposed method, *DL*_*n*_, achieved an agreement of 77.5%, 22.2%, and 8% for 1-peaks, 2-peaks, and 3-peaks, respectively, which was globally the closest to the agreement between the gold standards when compared to other methods. Although the single-fiber model (SFM) produced a relatively high level of agreement for 1-peaks with the ground truth (GT) at 84.6%, the agreement decreased to 4.6% and 2.5% for 2-peaks and 3-peaks, respectively. In contrast, the constrained spherical deconvolution (CSD) model achieved the lowest 1-peaks AR at 21.7%. This model showed a bias towards the estimation of multiple peaks, with 78% of the voxels modeled as having two or three peaks, which could be explained by the high b-value (b = 2600 *s/mm*^2^) that contains high levels of noise.

**Table 1.** Confusion matrices for number of peaks agreement (in %), normalized over all population. From left to right: gold standards GS1 vs. GS2, followed by the different methods CSA, CSD, SFM and *DL*_*n*_ compared to the GT MSMT-CSD. Each confusion matrix reports the average result for 320 test subjects (except SFM with 56 subjects).

| | $\Delta GS$ | | | CSD | | | CSA | | | SFM | | | $DL_n$ | | |
| --- | --- | --- | --- | --- | --- | --- | --- | --- | --- | --- | --- | --- | --- | --- | --- |
| #Fibers | 1 | 2 | 3 | 1 | 2 | 3 | 1 | 2 | 3 | 1 | 2 | 3 | 1 | 2 | 3 |
| 1 | 72.3 | 4.79 | 0.31 | 16.7 | 11.7 | 47.8 | 34.7 | 27.7 | 13.9 | 83.7 | 0.45 | 1.82 | 70.47 | 5.16 | 0.59 |
| 2 | 4.27 | 11.0 | 1.87 | 0.51 | 2.55 | 14.4 | 2.28 | 9.46 | 5.76 | 10.2 | 0.53 | 0.32 | 11.4 | 5.57 | 5.15 |
| 3 | 0.26 | 1.57 | 3.60 | 0.07 | 0.37 | 5.80 | 0.85 | 2.57 | 2.83 | 2.70 | 0.09 | 0.12 | 3.27 | 2.38 | 0.59 |

**Figure 3.**
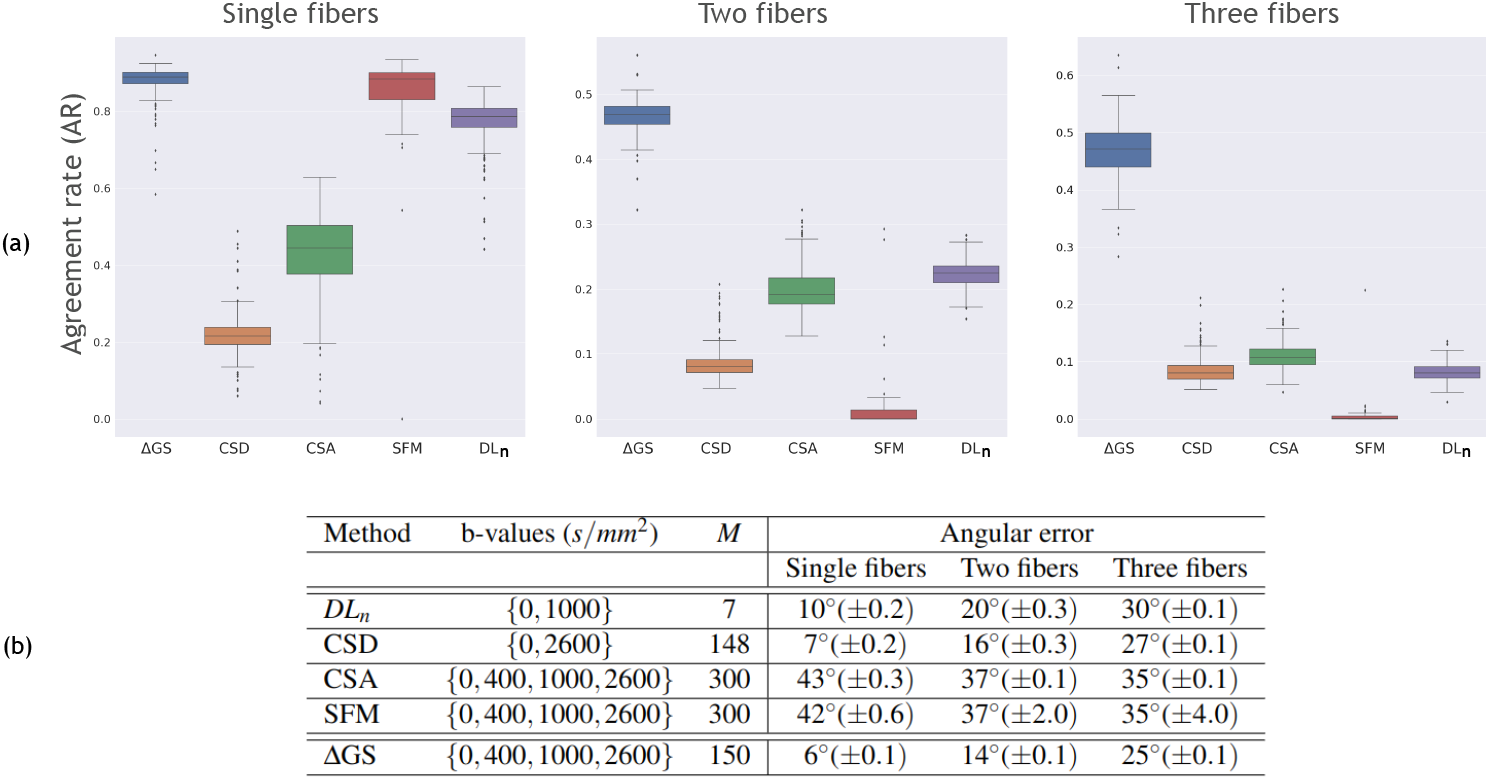
**(a)** Agreement rates, extracted from confusion matrices as defined in the Methods section, for different methods compared to the MSMT-CSD GT and for the agreement between the gold standard subsets. From right to left, the deep learning method using six measurements and *b0*, SFM and CSA using 300 multishell samples, CSD using 148 measurements and the agreement between the two gold standard (Δ*GS*) mutually exclusive subsets using each 150 samples. **(b)** Mean and standard deviation of angular error between GT (MSMT-CSD) and the different methods. ΔGS refers to GS1 and GS2 agreements. The number of measurements (*M*) and the b-values used are also reported. All results were statistically significant compared to ΔGS (*p* ≤9e^−10^). Our method achieves results comparable to the agreement rate ΔGS while using six measurements. It is worth noting that CSD is achieving slightly lower error because it misses more than 3 times GT-true single fiber voxels and more than two times GT-true two-fiber voxels, as can be seen in its low agreement rate in (a).

The relatively low agreement observed for voxels with multiple intravoxel fiber orientations might be attributed to their incongruence across the GT, resulting in the absence of a consistent pattern to be learned by the neural network. In fact, this is supported by the modest agreement between the two gold standards (Δ*GS*) where both the subjects and the number of measurements are the same, with only the gradient directions varying and already resulting in a drop of more than 50% in multiple fibers depiction. It is noteworthy that the agreement between different methods such as CSD versus CSA, SFM versus CSA, CSD versus *DL*_*n*_, among others, was also low. The confusion matrices for ΔGS agreement and the different methods can be found in Table 1.

Our analysis, presented in the table of Figure 3 (b), quantifies the angular error for different FOD methods. Our proposed learning model achieves an error rate that is comparable to GS1 and GS2. However, SFM and CSA methods demonstrate a higher error rate for single- and two-fiber voxels, whereas CSD outperforms the other techniques by achieving the lowest error rate. This could be attributed to the low AR observed for CSD, which reduces the error computation to a smaller subset of common voxels between the ground truth and CSD, as indicated in Figure 3 (a). The table of Figure 3 (b) also displays the number of measurements and the b-values that each method used. Notably, the angular error exhibits a nearly linear increase for voxels containing one, two, or three fibers. It is worth noting that training a network with 15 directions instead of 6 did not lead to a noticeable improvement in the results.

Finally, we explored the correlation with the quality control (QC) metrics provided by dHCP and the error metrics for Δ*GS*,

*DL*_*n*_, and CSD. The different error measures showed no correlation to QC metrics, i.e. SNR, outlier-ratio, nor scan age (Figure 10 in Supplementary Material for *DL*_*n*_) except for the motion as estimated by the SHARD^35^ pipeline. The more motion was estimated the higher the correlation to a lower agreement rate and a higher AFD error across subjects for the intra-agreement metrics of the MSMT-CSD GT (Δ*GS*) and both methods (*DL*_*n*_ and CSD) as can be shown in Figure 4. Statistical interaction analysis did not show however any significant difference in the way *DL*_*n*_ and CSD are influenced by motion (*p* = 0.8, *p* = 0.18 and *p* = 0.11 for single, two and three fibers respectively). Given that the motion was compensated^35^, we hypothesize that subjects with strong initial motion have still an increased residual motion after correction.

**Figure 4.**
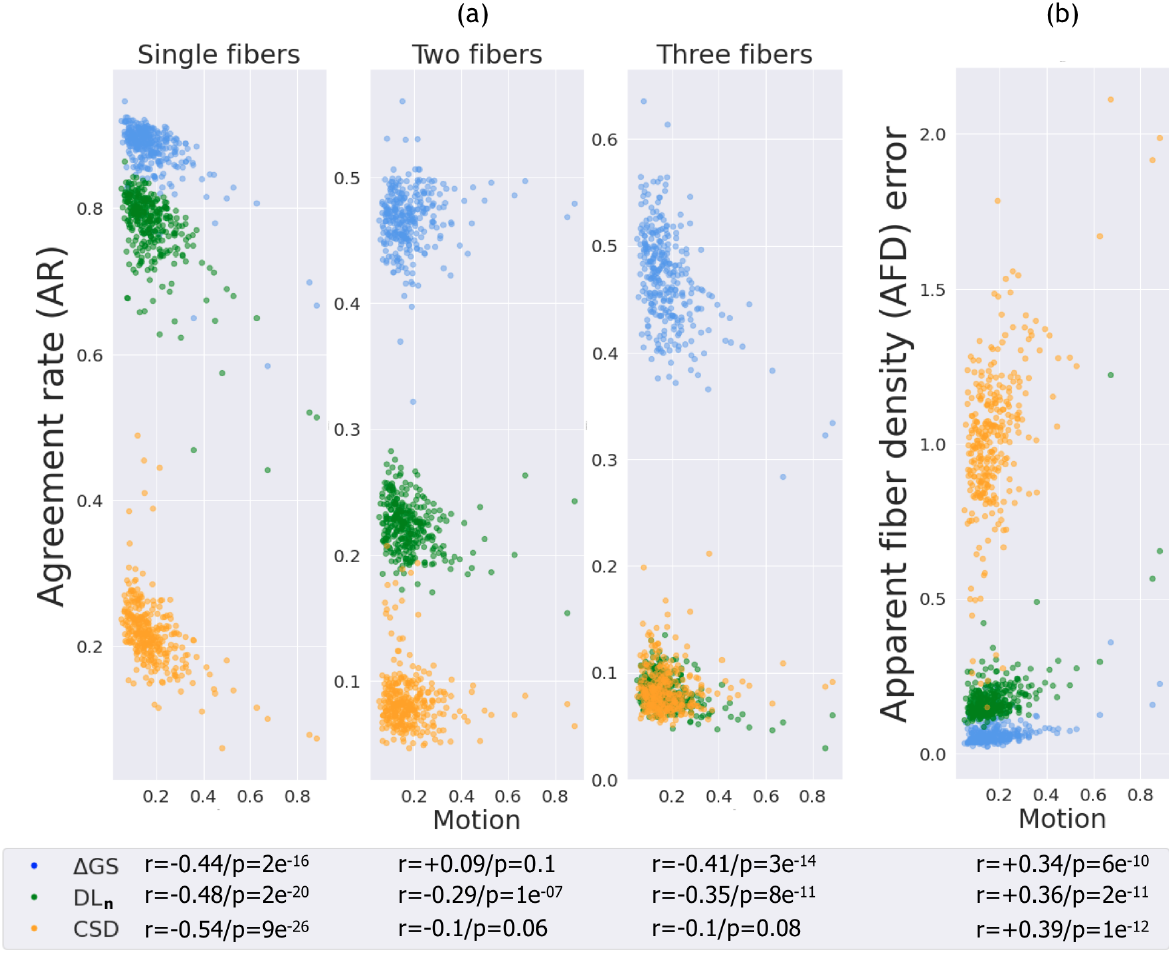
Agreement rate for voxels containing one, two, and three fibers, and apparent fiber density (AFD) error for the inter agreement between the gold standard datasets (Δ*GS*), deep learning method (*DL*_*n*_) and CSD, as a function of motion parameters (average translation and rotation parameters) for N=320 subjects. A negative correlation is generally observed with agreement rate (Spearman’s rank correlation coefficients shown in the figure with corresponding p-value). Similarly, a positive correlation with AFD error can be seen on the (b) panel. Other quality control (QC) metrics (Outlier ratio, Signal-to-noise ratio) and scan age didn’t exhibit any trend with the prediction accuracy (See Figure 10 in Supplementary Materials). Interaction analysis showed that CSD and *DL*_*n*_ were not significantly affected by motion (*p* ≥ 0.11).

#### Uncertainty

Using wild bootstrap (*N*_*WBS*_=60) on the six input directions of the 88 volumes of b = 1000 *s/mm*^2^ volumes, we have computed uncertainty maps using normalized standard deviation (please see Methods section). Figure 5 shows these maps compared to FA, where both images were applied a white matter mask. We can appreciate low uncertain regions in the highly anisotropic regions such as corpus callosum (body, splenium and genu) and internal capsules. In fact, this is in line with the results (Figure 3) where the *DL*_*n*_ model is less prone to errors in single coherent fiber populations as was previously studied in diffusion tensor imaging^36–38^. Hence, since these uncertainty maps do not need any ground truth and can express an increased correlation with erroneous predictions^39^, they can be used as an informative proxy to error detection, in case enough gradient directions are available for bootstrap.

**Figure 5.**
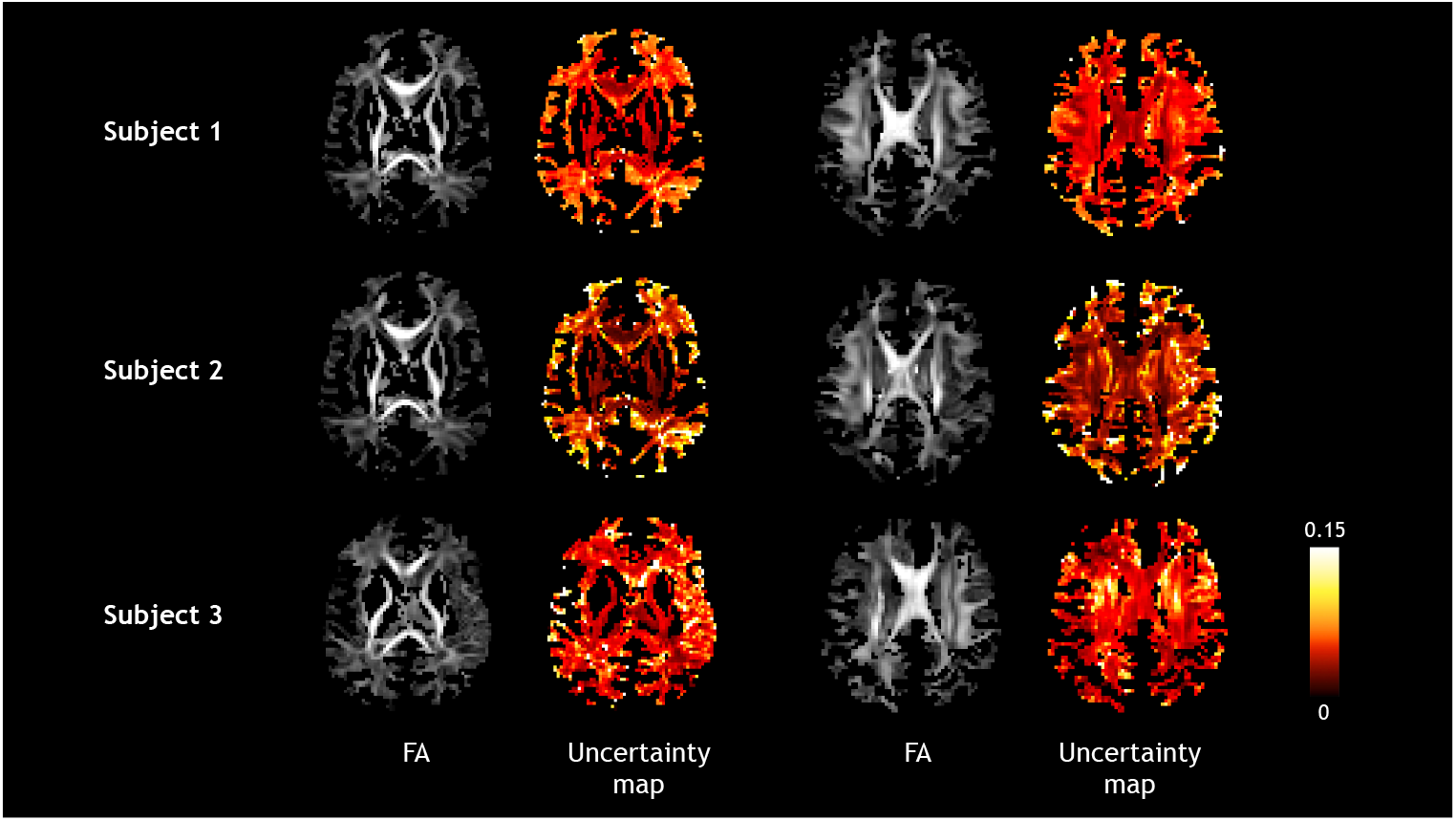
From top to bottom: three dHCP test subjects of respectively 43, 42 and 40 weeks. Uncertainty maps, computed using coefficient-normalized standard deviation of 60 bootstrapped gradient directions as described in the Methods section, are shown on the right. On the left, corresponding FA maps calculated from the diffusion tensor, that highlights regions of high anisotropy. Low uncertainty can be seen in such regions as the corpus callosum or the cortico-spinal tract, where the network has lower prediction errors. A white matter mask was applied to all images.

### Generalizability to clinical datasets

#### Neonates dMRI

The network *DL*_*n*_ trained on dHCP neonates was tested on 15 clinical newborns using six *b0*-normalized input volumes of b = 1000 *s/mm*^2^ as can be seen in Figure 6. As for the dHCP newborns, we can see the absence of high magnitude FODs in non-white matter regions, as opposite to noise-sensitive CSD (estimated using all 30 b = 1000 *s/mm*^2^ diffusion measurements and 5 *b0* volumes) that displays several false positive crossing fibers. These false crossings can also be noticed in some known single fiber areas such as the internal capsules as can be depicted in subject 1 of Figure 6.

**Figure 6.**
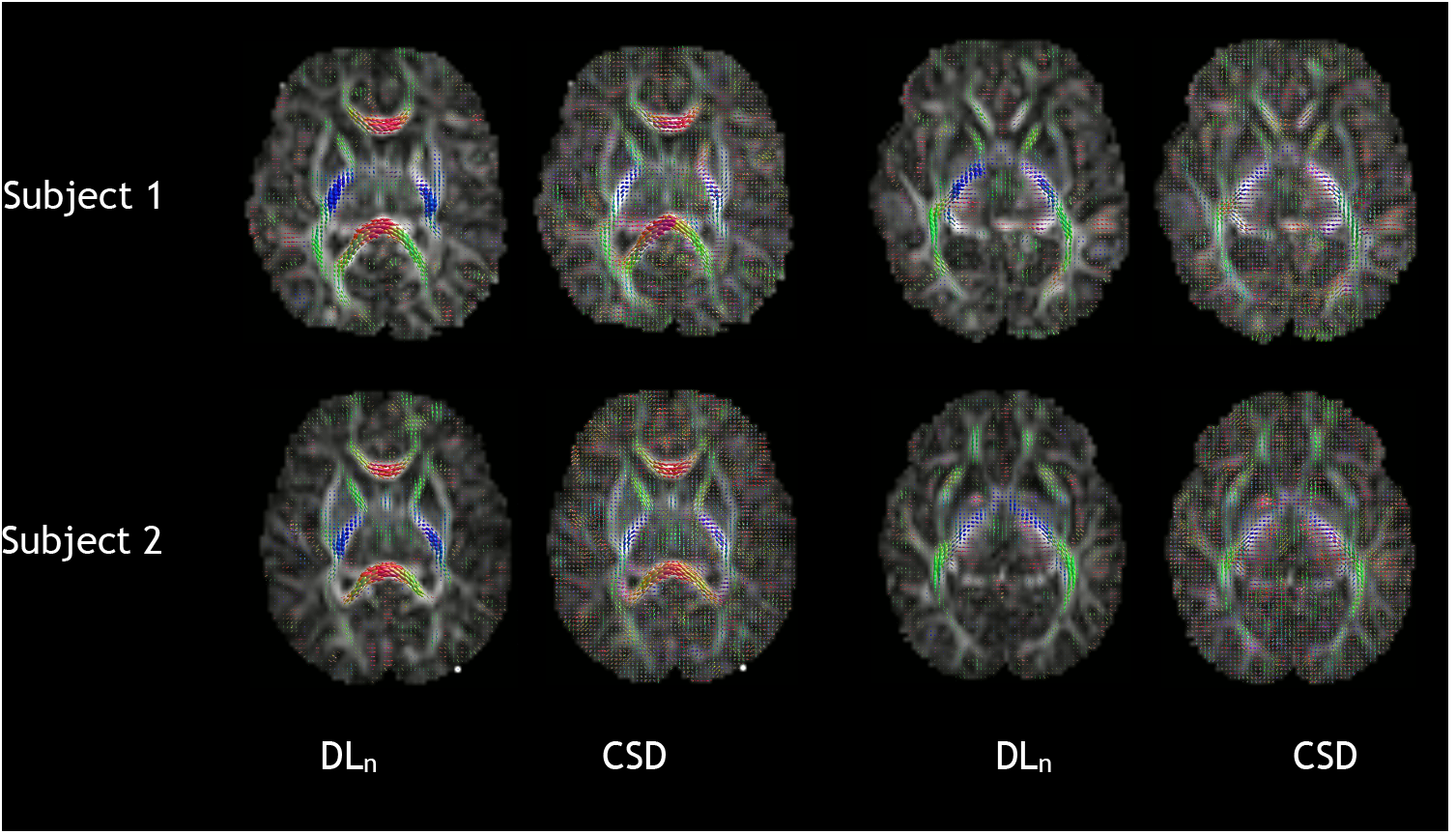
Qualitative comparison for two clinical newborn subjects (subject 1 and subject 2 of 41.8 and 38.1 weeks respectively) between the deep learning method *DL*_*n*_ (trained on dHCP dataset) using six b = 1000 *s/mm*^2^ measurements and one *b0*, and CSD using 30 measurements and 5 *b0* images. The background images are the corresponding fractional anisotropy (FA) maps.

#### In-utero fetal dMRI

We tested the proposed deep learning model, *DL*_*f*_, on 11 fetuses and compared it to CSD. In the absence of dMRI ground truth, we qualitatively evaluate the results. We have summarized the results for different anatomical regions in Table 2. We point out the frequency on which *DL*_*f*_ or CSD was depicted as better or in which they seem equivalent. The evaluation was conducted by an experienced developmental neuroanatomist and was based on former knowledge from histology.

**Table 2.** Comparison between the preferred method (*DL*_*f*_, CSD, or tied) for different regions of interest (ROI) in assessing the validity of the fibers in neurotypical fetal brains.

| Fetal brain region | $DL_f$ | Tied | CSD |
| --- | --- | --- | --- |
| Frontal crossroad region | 10 | 0 | 1 |
| Genu of corpus callosum | 11 | 0 | 0 |
| Cortex of Insula | 2 | 5 | 4 |
| Posterior limb of internal capsule | 7 | 2 | 2 |
| Cortex of superior temporal gyrus | 6 | 2 | 3 |
| Subplate of the precentral gyrus | 4 | 0 | 7 |
| Internal capsule | 3 | 0 | 8 |
| Cerebral peduncles | 1 | 1 | 9 |
| Intermediate zone, geniculocortical | 4 | 3 | 4 |
| Intermediate zones, callosal | 10 | 0 | 1 |
| Prefrontal subplate | 6 | 4 | 1 |
| Prefrontal cortical plate | 8 | 2 | 1 |
| **Count per ROI** | **7** | **1** | **3** |
| **Count per subject** | **9** | **0** | **2** |

This qualitative assessment relied on visually inspecting FOD maps in ROIs. We selected ROIs within regions whose tissue components are relatively known during early development. Specifically, these ROIs included: i) regions in the proximity of the frontal crossroad C2, ii) corpus callosum, iii) cortical plate (in the insula, superior temporal gyrus, prefrontal cortex), iv) subplate (precentral gyrus and sulcus and prefrontal cortex) v) internal capsule, vi) cerebral peduncles, and vii) intermediate zone (regions containing geniculocortical fibers, regions containing callosal fibers). Figure 7 depicts some of the aforementioned ROIs within two example subjects across the two methods on the corresponding FOD maps.

**Figure 7.**
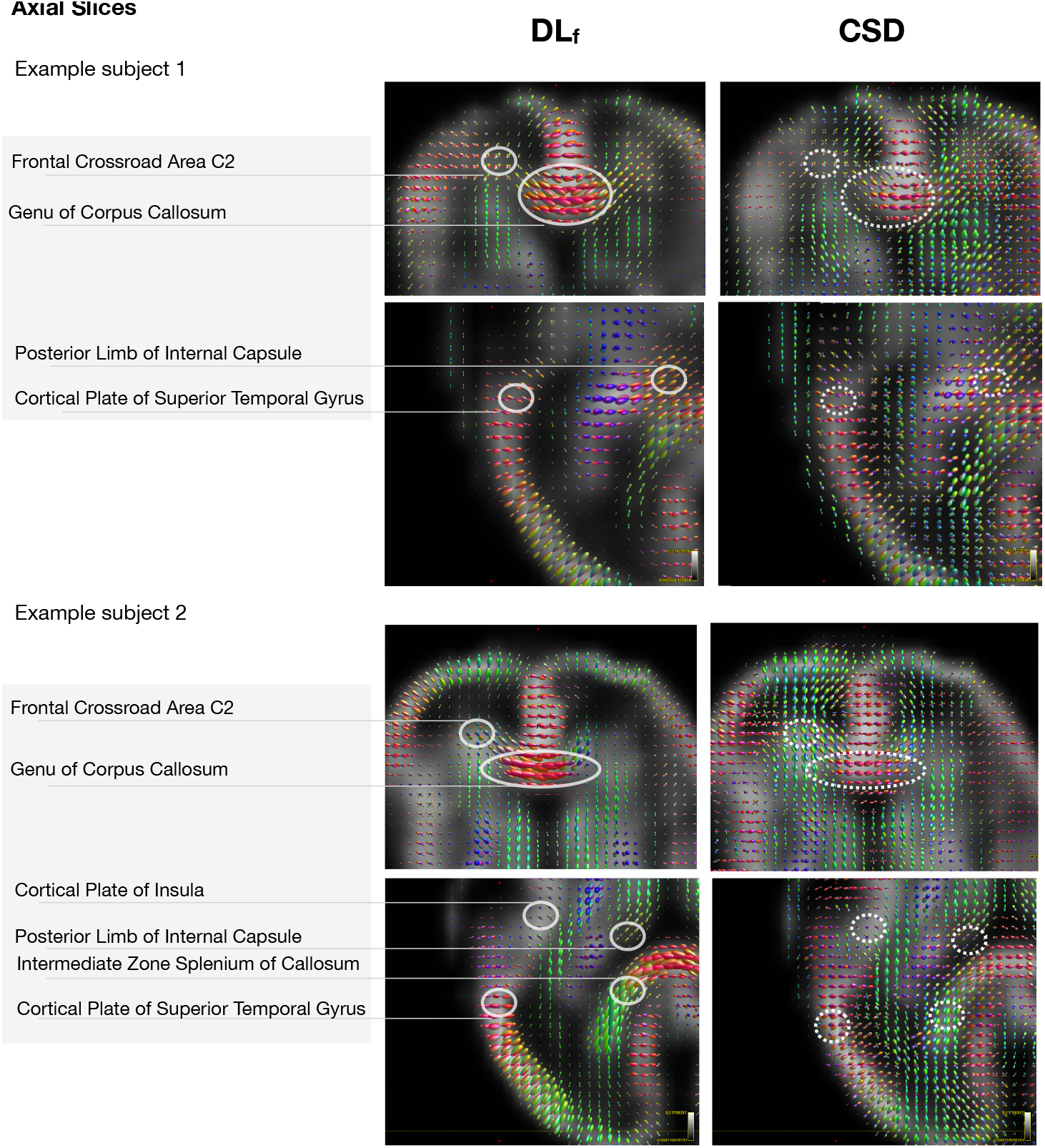
Qualitative assessment: Visual inspection of ROIs within FOD maps for two example subjects computed with *DL*_*f*_ and CSD. ROIs were selected based on the knowledge of microstructure from histology and immunohistochemistry.

Overall, *DL*_*f*_ performed better than CSD in predicting fiber orientation across most regions. Specifically, upon visual inspection, the regions surrounding the frontal crossroad region C2, genu of corpus callosum, intermediate zone containing callosal fibers, and prefrontal cortical plate were better defined using *DL*_*f*_. It is worth noting CSD systematically outperformed *DL*_*f*_ for cerebral peduncles and internal capsules on coronal sections. For the purpose of the current article, we added two slides to Figure 8 showing corresponding histology slices stained with GFAP (stains glial fibrillary acidic protein) or SMI 312 (stains highly phosphorylated axonal epitopes of neurofilaments)^40^. The criteria for evaluation included orientation, magnitude, and coherence of FODs. Specifically, in regions of corpus callosum^32^, cerebral peduncles, intermediate zone containing geniculocortical or callosal fibers, and internal capsule we expected high coherence with high magnitude, and orientation along or perpendicular to the main brain axes^32,41^. In contrast, within ROIs in the proximity of the frontal C2 crossroad^42^, we expected decreased coherence, with low magnitude and ambiguous orientation with FOD maps. Finally, the magnitudes, orientation, and coherence within the cortical plate and subplate ROIs were based on diffusion^30,41^ and histological descriptions of underlying microstructure^43^. As depicted in Figure 8, *DL*_*f*_ successfully defines the mediolateral orientation below the sulcus and rostrocaudal orientation of fibers in the gyrus.

**Figure 8.**
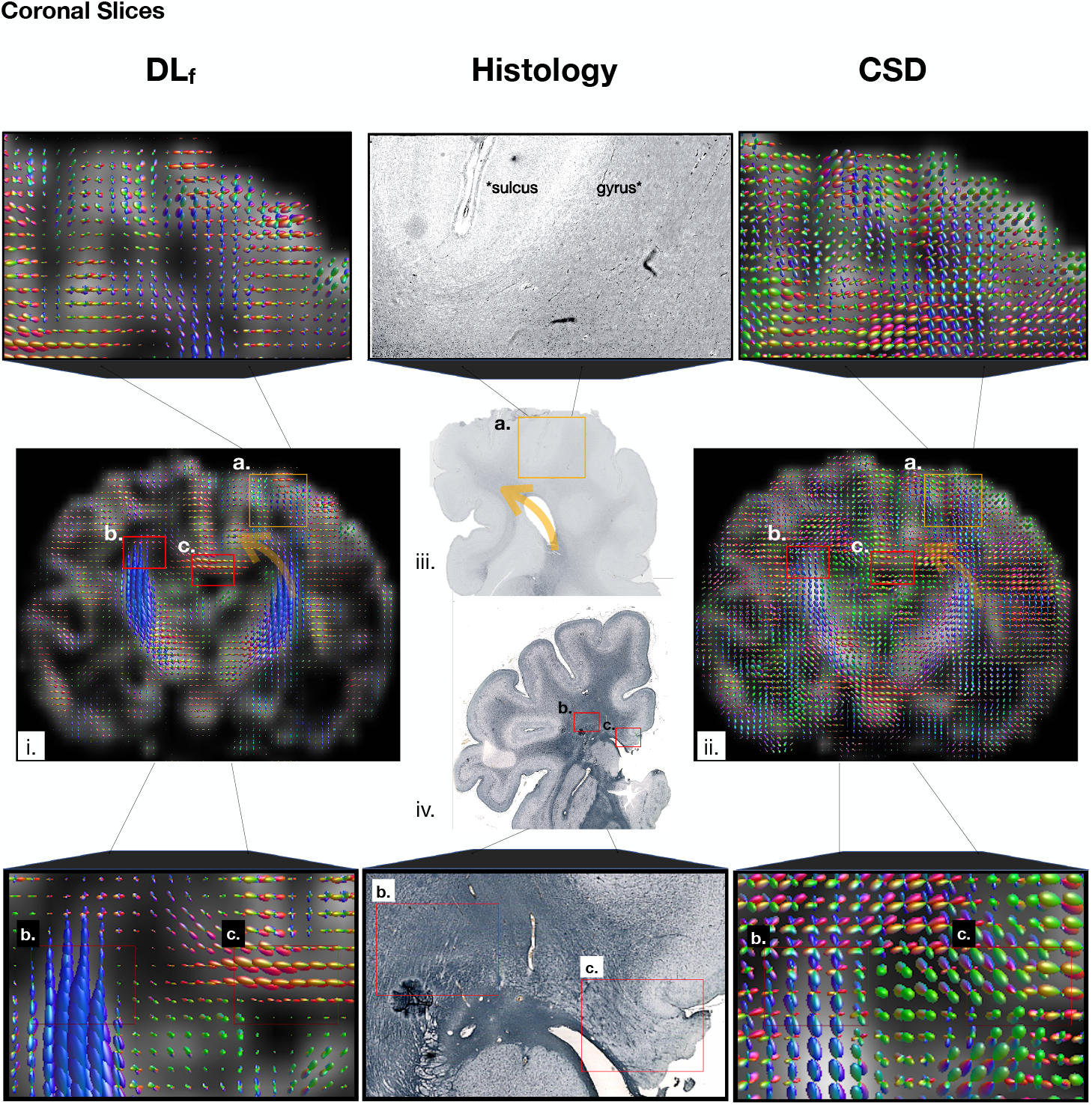
Qualitative assessment: Panel (iii.) shows a slice of a 40 GW fetal brain stained with SMI 312 directed against highly phosphorylated axonal epitopes of neurofilaments^40^, with rostral ROIs marked with an orange rectangle. This section is an example of coronal sections which were taken into consideration for the assessment of accuracy for our (i) *DL*_*f*_ and (ii) CSD method. Note the compactness of stained regions (marked with asterisks (*) in the magnified panels above figure iii. suggesting the mediolateral orientation of axonal fibers below the sulcus and rostrocaudal orientation with fanning of fibers within the gyrus. Corresponding regions are marked as (a.) within the FOD maps of both methods. Panel (iv.) shows another example of the coronal sections (40 GW fetal brain stained with GFAP) with 2 ROIs marked with red rectangles (e.g., the proximity of frontal crossroad area C2 (b.) and corpus callosum (c.)) that were taken into consideration for the assessment of accuracy for our (i) *DL*_*f*_ and (ii) CSD method. Note the compactness of GFAP-stained regions in red rectangles suggesting the orientation of axons in these regions in the magnified panels below.

## Discussion

In this study, we showed the effectiveness of deep neural networks in microstructure estimation of the developing human brain. Quantitative evaluation was performed on the highly controlled research oriented dHCP dataset where we show using several metrics that with six uniform measurements^44^, a carefully trained network can achieve performance level on par with the standard methods such as CSD. In particular, apparent fiber density^19^, a measure that is sensitive to fiber partial volume fraction was best estimated with the deep learning model.

We have additionally shown the out-of-domain generalization of the model to fetal and newborn clinical datasets, despite acquisition and anatomical gaps with respect to the training dataset. In fact, transferring knowledge from rich research oriented datasets (multiple b-values, multiple gradient directions, high magnetic field strength)^14,45^ to clinical datasets can be a winning strategy for developing cohorts such as from pre-terms to fetuses^28,46^ that cannot afford prolonged acquisition times because of increased motion, maternal discomfort and the sensitivity of this population. Moreover, to the best of our knowledge, this is the first diffusion MRI study to assess deep learning outputs with histology in the pediatric population, and the second across populations^47^.

Another important aspect of this study is the low agreement in estimating multiple intra-voxel fibers within the ground truth multi-shell multi-tissue CSD, hampering deep learning methods from learning consistent crossing fibers across different subjects. This highlights the need for acquisitions and reconstruction methods that are physically and anatomically informed, and tailored to the developing brain^48^. The data preprocessing has also an impact on the quality of the reconstruction. We have found an intra-agreement difference between two pipelines^35,49^ that reached 16% and 23% for the *agreement rate*, and 15° and 10°, for two and three fibers estimation^18^ respectively. Hence our simple and efficient strategy of splitting dMRI datasets into two independent subsets and computing their agreement for different metrics can provide a method to assess consistency of preprocessing pipelines. Lastly, we have shown that uncertainty maps computed using wild bootstrap can be a proxy for voxel-wise error detection.

Recently, fiber orientation distribution function prediction using deep learning has received growing interest spanning several goals that go beyond the objective of directly learning FODs from raw data or its spherical harmonics representation^27–29^. For instance a recent study aimed at mapping 3T dMRI data to 7T FODs^50^, while another work simultaneously learned FODs from all radial combinations of multi-shell data using spherical convolutions^51^. This last study can be particularly interesting to increase the generalizability of our work to multiple b-values. In fact, our model generalization from pre-term to fetuses is also due to the proximity of the two acquired b-values for training and evaluation (b = 400 *s/mm*^2^ and b = 500 *s/mm*^2^, respectively). In contrast, our network has failed predicting coherent FODs of a different fetal dataset acquired at b = 700 *s/mm*^2^, likely because of the lower SNR and contrast difference, despite training our network on *b0*-normalized images. This limitation can partly be explained by specific response functions that were learned by the network. Hence, strategies enhancing generalization^51^ and data harmonization^52^ can be adapted for future work.

Another limitation of this study is the absence of pathological datasets, which we aim to include in future work. We also intend to incorporate in the neural network, convolutions and deconvolutions that take into account the spherical property of the diffusion signal (angular dimension) such as roto-translation equivariant convolutions^53^. Moreover, as there is no consensus on the diffusion protocol of fetal brain diffusion imaging^45,54–57^, we want to explore optimal gradient tables that recover the most accurate white matter representation.

## Methods

### Data

#### Research dMRI acquisition protocol in neonates

We used the data from the third release of the publicly available dHCP dataset ^1^. Scans were performed on a 3T Philips Achieva system with a customized 32-channel neonatal head coil. The protocol employed a TE of 90ms, TR of 3800ms, a multiband factor of 4, a SENSE factor of 1.2, a Partial Fourier factor of 0.855, a 1.5mm in-plane resolution, and 3mm slice thickness with 1.5mm slice overlap^14^. The diffusion gradient scheme used four shells {0, 400, 1000, 2600} s/mm^2^ with 20, 64, 88, and 128 samples, respectively. The slice order was interleaved with a factor of 3 and a shift of 2. Data were processed and reconstructed with the SHARD^35,58^ pipeline that included denoising, Gibbs ringing suppression, distortion correction and motion correction. The resolution of the processed data is 1.5 mm^3^ isotropic with a field of view of 100 × 100 × 64 voxels.

Two subsets were extracted from the SHARD-processed dataset, (i) 465 subjects with postmenstrual age range [26.71, 45.14] weeks (mean±std = 39.75±3.05 weeks), and (ii) a group of 77 pre-term subjects with postmenstrual ages ranging from 26.71 to 38.0 weeks (mean±std = 34.79±2.52 weeks). We generated a white matter mask by combining the *White Matter* and the *Brainstem* labels provided by the dHCP with the voxels where Fractional Anisotropy (FA) was greater than 0.25. Finally, the dHCP labels were resampled from T2-w resolution (0.5 mm^3^ isotropic) to 1.5 mm^3^ resolution.

#### Clinical dMRI acquisitions in neonates

We retrospectively used data from 15 newborns at a postmenstrual age between [38.14, 48] weeks (mean±std=41.25±2.34 weeks), while they were in natural sleep and scanned using Siemens Trio or Skyra MRI machines at 3T. The imaging protocol included acquiring five *b0* images and 30 diffusion-weighted images with b = 1000 s/mm^2^. The TR-TE values used were 3700-104 ms, and the voxel size was 2 mm isotropic. The images were resampled to 1.5 mm^3^ isotropic resolution.

#### Clinical dMRI acquisitions in fetuses

A total of 11 fetuses each scanned at a gestational age between [24, 38.71] weeks (mean±std=28.89±4.6 GW) were included in this study. All subjects were scanned using a 3T Siemens Skyra MRI scanner, with one *b0* and 12 diffusion-sensitized images at b = 500 *s/mm*^2^, with a TR of 3000–4000 ms and a TE of 60 ms. The diffusion scans were repeated between 2 to 5 times during acquisitions, and one of the scans with the least amount of fetal motion was chosen for the analysis. Preprocessing of the data was performed to correct for noise^59^ and bias field inhomogeneities^60^. Registration of the images to a T2 atlas^61^ was carried out using rigid transformation, and b-vectors were subsequently rotated accordingly. The resulting images were upsampled from 2 × 2 × 3 − 4 mm^3^ to 1.5 mm^3^. Ethical approval for this study on neonatal and fetal MRI scans was granted by the Institutional Review Board Committee at Boston Children’s Hospital.

#### Histological post-mortem fetuses

Neonatal post-mortem human brain specimens without evident pathological changes are part the Zagreb Collection of Human Brains^62^. Tissue was obtained during regular autopsies either after spontaneous abortions, or after the death of prematurely born infants at the clinical hospitals associated to the University of Zagreb, School of Medicine. After fixation in 4% paraformaldehyde (PFA), tissue was embedded in paraffin. Sections were cut in coronal plane and proceeded with routine immunohistochemistry protocol. In brief, after deparaffinization and 0.3% hydrogen peroxide treatment, sections were incubated in blocking solution: 3% bovine serum albumin BSA and 0.5% Triton x-100 (Sigma, St. Louis, MO) in 0.1M PBS. Next, sections were incubated with primary antibodies (anti-GFAP, Dako, z-0334, 1:1000; anti-SMI-312 [panaxonal anti-neurofilament marker], Biolegend, 837904, 1:1000) at room temperature overnight. Following washes, sections were incubated with secondary, biotinylated antibodies according to manufacturer’s protocol (Vectastain ABC kit, Vector Laboratories, Burlingame, CA). Staining was developed using 3,3-diaminobenzidine (DAB) with enhancer (Sigma, St. Louis, MO) and slides were coverslipped (Histomount mounting medium, National Diagnostics, Charlotte, NC). Finally, staining was visualised by a high-resolution digital slide scanner NanoZoomer 2.0RS (Hamamatsu, Japan). Tissue sampling was performed in agreement with the Declaration of Helsinki, 2000, previously approved by the Institutional Review Board of the Ethical Committee, University of Zagreb, School of Medicine.

### Model

Our study employed two different neural networks, one for inference on neonates (*DL*_*n*_) and another for inference on fetuses (*DL*_*f*_). *DL*_*n*_ was trained on newborn subjects using six single-shell (b = 1000 *s/mm*^2^) measurements, and *DL*_*f*_ was trained on pre-term subjects using twelve single-shell (b = 400 *s/mm*^2^) measurements. To make the model independent on gradient directions, we projected the signal onto spherical harmonics basis (SH) with SH-*L*_*max*_ order 2 and 4 for *DL*_*n*_ and *DL*_*f*_, respectively, to predict the fiber orientation distribution (FOD) represented in the SH basis with SH-*L*_*max*_ order 8. The latter is composed of 45 coefficients (45 channels for the network) and was generated using 300 multi-shell measurements obtained using MSMT-CSD^18^. These measurements were distributed over three shells with b-values of 400, 1000, 2600 s/mm^2^ and had 64, 88, and 128 samples, respectively, along with 20 *b0* (b = 0 *s/mm*^2^) images. The input measurements for the model were based on the scheme proposed by Skare et al.^44^, which minimized the *condition number* of the diffusion tensor reconstruction matrix.

### Network architecture

The deep convolutional neural network can be seen in the yellow box of Figure 9. It aims at directly learning a cascade of features (i.e. *feature maps*) from the input data that can predict the target FODs, namely the 45 SH coefficients. These features are randomly selected in the beginning and are gradually learned, after each iteration, by minimizing the error between the ground truth FODs and the predicted FODs by the network. The network architecture resembles that of U-Net^63^ with two main modifications. Firstly, the network has extensive short and long-range residual connections, which provide more context to subsequent layers. This design choice is particularly important given the low dimensionality of our input (6 channels) compared to the output (45 channels). Secondly, the conventional max-pooling operations in the contracting path have been replaced with stride-2 convolutions to enable downsampling as a learnable step that is specific to each layer. The first block is set to 36 feature maps that are doubled after each contracting block. Each layer is composed of convolutions that are followed by Rectified Linear Unit (ReLu)^64^ activation functions, followed by a dropout^65^ layer. For the output layer, neither ReLu nor dropout were applied.

**Figure 9.**
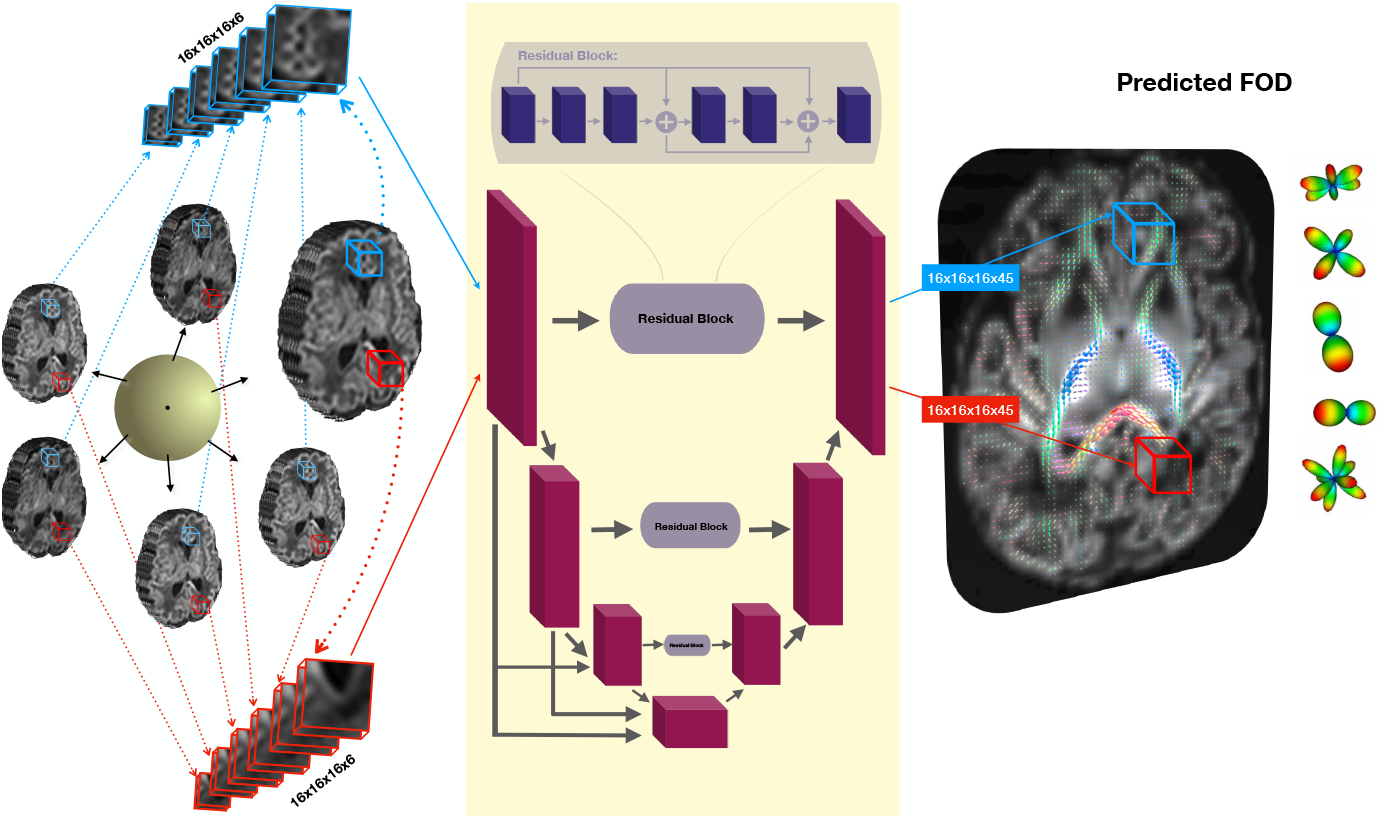
Schematic illustration of the proposed deep learning framework for predicting the Fiber Orientation Distribution (FOD). The input to the network consists of 3D patches derived from 6 diffusion measurements, which are normalized with *b0*. The network predicts the spherical harmonic coefficients (of order SH-*L*_*max*_ = 8) of the FOD for the input patch. Two example patches are shown in blue and red. The network trained on pre-term newborns takes 12 instead of 6 measurements.

**Figure 10.**
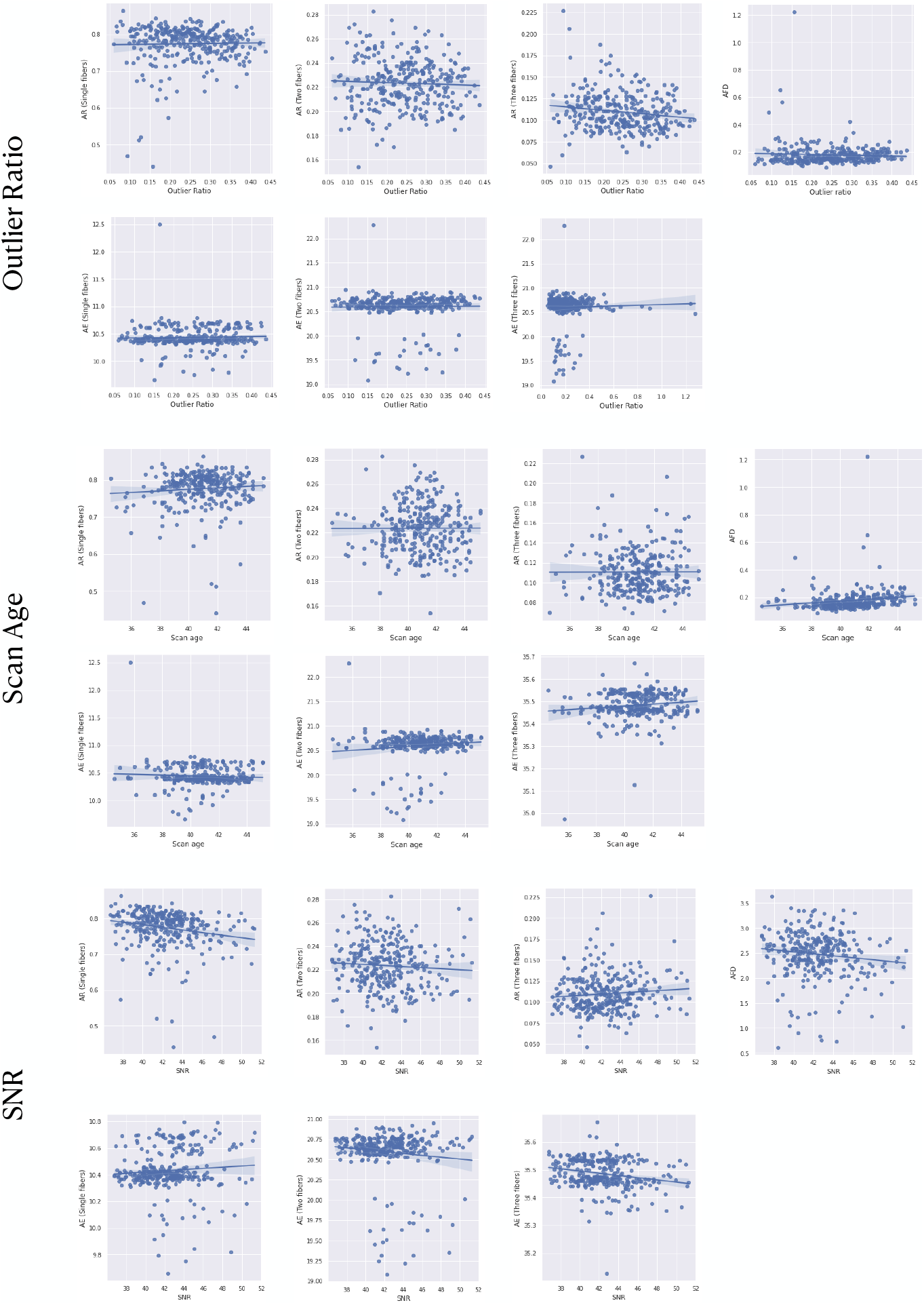
Agreement rate (AR) and angular error (AE) for single, two and three fibers, and apparent fiber density (AFD) error for the deep learning method (*DL*_*n*_), as a function of quality control (QC) metrics (Outlier ratio, Signal-to-noise ratio) and scan age for N=320 subjects. No correlation is generally observed between the QC metrics and the error rates.

### Training strategies

The input data was first normalized by *b0* to improve network convergence and to reduce b-value dependency. Data were split into training, validation and test sets: for *DL*_*n*_, 109, 36 and 320 subjects, respectively; for *DL*_*f*_, 58, 19, pre-terms and 11 fetal subjects, respectively. To ensure balanced patch selection per batch, the number of FOD peaks (extracted using Dipy^66^) was used as a criterion. The central voxel of each patch was restricted to be in the generated white matter mask and to have one peak in 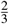 of the batch and more than one peak in 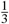 of the batch. This condition ensured that empty patches were not selected. The patch size was empirically varied in {8^3^, 16^3^, 36^3^, 48^3^} voxels (no performance increase was observed for patches bigger than 16^3^, so all networks used 16^3^ voxels). At testing, the method employed a sliding window technique to sequentially process all non-empty patches.

We used Adam optimizer^67^ to minimize the 𝓁_2_ norm loss function between the predicted 45 spherical harmonic (SH) coefficients and the ground truth fiber orientation distribution (FOD) SH coefficients. Since the order of the magnitude of the coefficients depends on which SH-order the coefficient belongs to, we have used pre-defined weights to penalize small coefficients. These weights were inversely proportional to the order of the magnitude of the coefficient in the GT. Namely, these weights were proportional to the first SH coefficient and were around 2.5, 4, 7.5 and 20 for coefficients of SH order 2, 4, 6 and 8, respectively. However, no gain was observed with this scheme so all coefficient weights were set to 1.

The batch size was set to 27 for *DL*_*n*_ and 9 for *DL*_*f*_, and the initial learning rate was set to 10^−4^. The learning rate was decreased by a factor of 0.9 whenever the validation loss did not improve after one epoch. The total number of training epochs was 10000, and a dropout rate of 0.1 was used in all layers to reduce overfitting and improve generalization. In *DL*_*f*_, Gaussian noise (*μ* = 0, *σ* = 0.025) was injected to the input training data to encourage robustness to noise in fetal data. Moreover, small rotations (uniformly from [−5°, +5°]) were applied to improve the robustness of the model to minor uncorrected movements due to small differences in scanning field of view and fetal head motion.

### Implementation details

All models were implemented using TensorFlow (1.6) and run on an NVIDIA GeForce GTX A6000 on a Linux machine with 125 GB of memory and 20 CPU cores. Convergence of each model took approximately 40 hours. Testing takes less than 1 minute per subject on the same machine. Code will be made publicly available.

### Evaluation

Quantitative evaluation has been carried out for *DL*_*n*_ predictions compared to the GT MSMT-CSD. Moreover, three state-of-the-art techniques were computed as baseline models, namely: Constrained Spherical Deconvolution (CSD) method^17^, using 128 gradient directions from the highest shell (b-value of 2600 *s/mm*^2^) and 20 *b0* images; Constant Solid Angle model^33^ which is referred to as CSA; and the Sparse Fascicle Model (SFM)^34^ for which the default regularization parameters were employed. The latter was only applied on 57 dHCP subjects as it takes a significant time to be run (around 7 hours per subject).

*DL*_*n*_, using 6 *b0*-normalized diffusion measurements, was also tested on the clinical neonate dataset and compared against CSD using all measurements (35). Similarly, *DL*_*f*_ using 12 *b0*-normalized diffusion samples was tested and evaluated against CSD. Because clinical datasets do not have densely sampled (multiple-shell) measurements that can be considered as high quality ground truth, only qualitative evaluation was performed.

#### Agreement within ground truth (ΔGS)

We applied MSMT-CSD on two mutually exclusive subsets of the 300 measurements from the dHCP dataset. We denote the estimation results from these two subsets with GS1 and GS2, and the difference between these with Δ*GS*. Each subgroup consists of 150 samples at b ∈ 0, 400, 1000, 2600 s/mm^2^ with 10, 32, 44, and 64 scans respectively. Both GS1 and GS2 can be regarded as independent high-quality scans of the same subject with a similar protocol, i.e., the same b-values and the same number but different gradient directions. Hence, any discrepancies between them with respect to diffusion metrics can be considered as an error upper bound of errors between the different methods and the full GT.

#### Quantitative performance on dHCP dataset

We conducted a quantitative validation by evaluating the performance of different fiber orientation distribution (FOD) estimation methods. The validation was based on three metrics, namely, the **number of peaks, angular error, and the apparent fiber density (AFD)**^19^. The number of peaks was computed for the FODs predicted by the network and those estimated by various methods (GT, GS1, GS2, CSD, CSA, and SFM). We set up a maximum number of 3 peaks, a mean separation angle of 45°, and a relative peak threshold of 0.5. The choice of these parameters was guided by the work of Schilling et al.^68^, which demonstrated the limitations of current diffusion MRI models in correctly estimating multiple fiber populations and low angular crossing fibers. We compared the different models based on confusion matrices and the **agreement rate (AR)** that is extracted from the latter. AR was defined for each number of peaks *p* as:

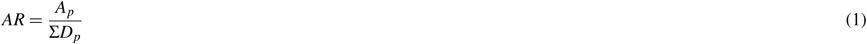

where *A*_*p*_ represents the percentage of voxels where both methods agree on *p* number of peaks and *D*_*p*_ denotes the percentage of voxels where at least one of the two methods predicts *p* and the other predicts *p*^′^ (*p* ≠ *p*^′^). This metric hence captures intuitively the rate of concordance between two methods.

**Mean angular error** was also computed for voxels containing the same number of estimated peaks. For voxels with multiple fibers, we extracted the corresponding peaks between the selected method and the GT (or the agreement between GS1 and GS) by computing the minimum angle between all configurations, namely 4 configurations for 2 peaks and 9 for 3 peaks. We subsequently eliminated those peaks and applied the same algorithm recursively until all peaks are matched. We also compared the error related to the apparent fiber density (FOD amplitude) along with the agreement between GS1 and GS2. We performed a statistical validation using paired t-test corrected for multiple comparisons with Bonferroni method to compare the errors of the different methods with respect to GT and the difference between GS1 and GS2.

The different error measures were correlated to quality control (QC) metrics provided by the SHARD pipeline of dHCP^35^. Namely, Signal-to-Noise Ratio that is calculated from denoising residuals; (2) Motion metrics, i.e. translation and rotation quantifying subject activity during scan and (3) Outlier ratio, as detected in slice-to-volume reconstruction^35^. We averaged both translation and rotation metrics to have one metric that we label as *motion*. We have also added the age of scan to the different QC metrics to check for any potential correlation. We have performed this analysis for the GT MSMT-CSD to assess the consistency of the dataset across the QC metrics.

#### Qualitative assessment of clinical datasets

Detailed assessment was performed for the FODs generated on the clinical fetal dataset by an expert fetal neuroanatomist (LV). The images were blinded and the method used to reconstruct the maps was masked for the reader. The 12 ROIs were selected based on the anatomical knowledge (previously reported in Kunz et al. 2014^69^). Next, the corresponding slices of volumes reconstructed with both methods were placed side by side and FODs were inspected in each ROIs using MRView^70^. Based on the visual inspection and taking into the consideration coherence, orientation, and magnitudes, the ROIs were marked as ‘better with *DL*_*f*_’, ‘better with CSD’, or equal. After examining all the brains, we generated the table and counted ROIs and subjects where *DL*_*f*_ outperformed CSD, CSD outperformed *DL*_*f*_ or tied.

#### Uncertainty estimation

A metric that expresses an increased likelihood on erroneous predictions can be very valuable in the absence of ground truth. Uncertainty in that sense can be used for that aim. Post-hoc uncertainty using wild bootstrap that has been used in diffusion tensor imaging^71–73^ was the method of choice that was most suited to our study. We randomly selected 6 gradient directions (*N*_*WBS*_=60) from the 88 samples of the b = 1000 *s/mm*^2^ shell from the dHCP data. The 6 directions were constrained to have a *condition number*^44^ of at most 2 to guarantee that the b-vectors are minimally uniformly distributed. We then computed for each voxel, the standard deviation of the predicted FODs of the *N*_*WBS*_ bootstrapped volumes (Equation 2). Given that FOD coefficients have different orders of magnitude, this standard deviation was normalized by the norm of the FOD (Equation 3). Specifically, for each voxel we calculate *σ*_*norm*_ that we define as our uncertainty measure from:

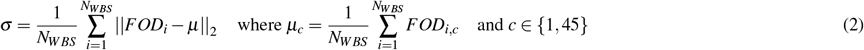

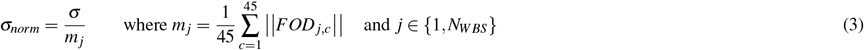

## Acknowledgements

This work was supported by the Swiss National Science Foundation (project 205321-182602). We acknowledge access to the facilities and expertise of the CIBM Center for Biomedical Imaging, a Swiss research center of excellence founded and supported by Lausanne University Hospital (CHUV), University of Lausanne (UNIL), Ecole polytechnique fédérale de Lausanne (EPFL), University of Geneva (UNIGE) and Geneva University Hospitals (HUG).

This research was supported in part by the National Institute of Biomedical Imaging and Bioengineering, the National Institute of Neurological Disorders and Stroke, and Eunice Kennedy Shriver National Institute of Child Health and Human Development of the National Institutes of Health (NIH) under award numbers R01HD110772, R01NS128281, R01NS106030, R01EB031849, R01EB032366, and R01HD109395; in part by the Office of the Director of the NIH under award number S10OD025111; and in part by the National Science Foundation (NSF) under grant number 212306. This research was also partly supported by NVIDIA Corporation and utilized NVIDIA RTX A6000 and RTX A5000 GPUs. The content of this publication is solely the responsibility of the authors and does not necessarily represent the official views of the NIH, NSF, or NVIDIA.

We would like to thank Dr. Maria Deprez and Dr. Daan Christiaens from King’s College London (London, United Kingdom) for their support in answering questions about the dHCP dataset. We would also like to thank Dr. Farnaz Delavari from University of Geneva (Geneva, Switzerland) for her help in the design of the figures and statistical analysis.

## Author contributions statement

HK performed the technical analysis, wrote the manuscript and contributed to the conceptualization of the research project. DK provided the original idea and the original code. LV evaluated the clinical datasets and contributed to the manuscript. ZK provided the histological dataset. DK, AG, and MBC conceptualized, designed and supervised the research project, contributed to the manuscript. MBC provided funding and AG provided infrastructure in the host lab of HK. All authors contributed to the article and approved the submitted version.

## Ethics declarations

Ethical approval for this study on neonatal and fetal MRI scans was granted by the Institutional Review Board Committee at Boston Children’s Hospital. Newborn subject scans that were acquired as a part of the dHCP were approved by the National Research Ethics Committee and informed written consent given by the parents of all participants.

## Data availability

Part of the analyzed datasets were publicly available. This data can be found in https://www.developingconnectome.org/data-release/third-data-release/

## Code availability

The code will be made publicly available in https://github.com/medical-Image-Analysis-Laboratory/

## Supplementary Material

## Footnotes

1 https://www.developingconnectome.org/data-release/third-data-release/

## References

1. Volpe, J. Brain injury in premature infants: A complex amalgam of destructive and developmental disturbances. Lancet Neurol. 8, 110–124, DOI: 10.1016/S1474-4422(08)70294-1 (2009).

2. Bhat, S., Acharya, U., Adeli, H., Bairy, G. & Adeli, A. Autism: Cause factors, early diagnosis and therapies. Rev. Neurosci. 25, 841–850, DOI: 10.1515/revneuro-2014-0056 (2014).

3. Kwon, E. & Kim, Y. What is fetal programming?: A lifetime health is under the control of in utero health. Obstet. & Gynecol. Sci. 60, 506–519, DOI: 10.5468/ogs.2017.60.6.506 (2017).

4. O’Donnell, K. & Meaney, M. Fetal origins of mental health: The developmental origins of health and disease hypothesis. The Am. J. Psychiatry 174, 319–328, DOI: 10.1176/appi.ajp.2016.16020138 (2017).

5. Bilder, D. et al. Early second trimester maternal serum steroid-related biomarkers associated with autism spectrum disorder. J. autism developmental disorders 49, 4572–4583, DOI: 10.1007/s10803-019-04162-2 (2019).

6. Stejskal, E. O. & Tanner, J. E. Spin diffusion measurements: spin echoes in the presence of a time-dependent field gradient. The journal chemical physics 42, 288–292 (1965).

7. Le Bihan, D. et al. Mr imaging of intravoxel incoherent motions: application to diffusion and perfusion in neurologic disorders. Radiology 161, 401–407 (1986).

8. Dubois, J. et al. The early development of brain white matter: a review of imaging studies in fetuses, newborns and infants. neuroscience 276, 48–71 (2014).

9. Christiaens, D. et al. In utero diffusion mri: challenges, advances, and applications. Top. Magn. Reson. Imaging 28, 255–264 (2019).

10. Jakab, A. et al. Fetal cerebral magnetic resonance imaging beyond morphology. In Seminars in Ultrasound, CT and MRI, vol. 36, 465–475 (Elsevier, 2015).

11. Wilson, S. et al. Development of human white matter pathways in utero over the second and third trimester. Proc. Natl. Acad. Sci. 118, e2023598118 (2021).

12. Xu, X. et al. Spatiotemporal atlas of the fetal brain depicts cortical developmental gradient. J. Neurosci. 42, 9435–9449 (2022).

13. Calixto, C. et al. Detailed anatomic segmentations of a fetal brain diffusion tensor imaging atlas between 23 and 30 weeks of gestation. Hum. Brain Mapp. 44, 1593–1602 (2023).

14. Hutter, J. et al. Time-efficient and flexible design of optimized multishell hardi diffusion. Magn. resonance medicine 79, 1276–1292 (2018).

15. Tournier, J.-D. et al. A data-driven approach to optimising the encoding for multi-shell diffusion mri with application to neonatal imaging. NMR Biomed. 33, e4348 (2020).

16. Basser, P. J., Mattiello, J. & LeBihan, D. Mr diffusion tensor spectroscopy and imaging. Biophys. journal 66, 259–267 (1994).

17. Tournier, J.-D. et al. Direct estimation of the fiber orientation density function from diffusion-weighted mri data using spherical deconvolution. Neuroimage 23, 1176–1185 (2004).

18. Jeurissen, B. et al. Multi-tissue constrained spherical deconvolution for improved analysis of multi-shell diffusion mri data. Neuroimage 103, 411–426 (2014).

19. Raffelt, D. et al. Apparent fibre density: a novel measure for the analysis of diffusion-weighted magnetic resonance images. Neuroimage 59, 3976–3994 (2012).

20. Jbabdi, S. & Johansen-Berg, H. Tractography: where do we go from here? Brain connectivity 1, 169–183 (2011).

21. Jeurissen, B. et al. Diffusion mri fiber tractography of the brain. NMR Biomed. 32, e3785 (2019).

22. Golkov, V. et al. Q-space deep learning: twelve-fold shorter and model-free diffusion mri scans. IEEE TMI 35, 1344–1351 (2016).

23. Lu, H., Jensen, J. H., Ramani, A. & Helpern, J. A. Three-dimensional characterization of non-gaussian water diffusion in humans using diffusion kurtosis imaging. NMR Biomed. An Int. J. Devoted to Dev. Appl. Magn. Reson. In vivo 19, 236–247 (2006).

24. Zhang, H., Schneider, T., Wheeler-Kingshott, C. A. & Alexander, D. C. Noddi: practical in vivo neurite orientation dispersion and density imaging of the human brain. Neuroimage 61, 1000–1016 (2012).

25. Li, H. et al. Superdti: Ultrafast dti and fiber tractography with deep learning. Magn. resonance medicine 86, 3334–3347 (2021).

26. Koppers, S. & Merhof, D. Direct estimation of fiber orientations using deep learning in diffusion imaging. In International Workshop on Machine Learning in Medical Imaging, 53–60 (Springer, 2016).

27. Lin, Z. et al. Fast learning of fiber orientation distribution function for mr tractography using convolutional neural network. Med. physics 46, 3101–3116 (2019).

28. Karimi, D. et al. Deep learning-based parameter estimation in fetal diffusion-weighted mri. Neuroimage 243, 118482 (2021).

29. Hosseini, S. et al. Cttrack: A cnn+ transformer-based framework for fiber orientation estimation & tractography. Neurosci. Informatics 2, 100099 (2022).

30. Khan, S. et al. Fetal brain growth portrayed by a spatiotemporal diffusion tensor mri atlas computed from in utero images. Neuroimage 185, 593–608 (2019).

31. Huang, H. & Vasung, L. Gaining insight of fetal brain development with diffusion mri and histology. Int. J. Dev. Neurosci. 32, 11–22 (2014).

32. Vasung, L., Raguz, M., Kostovic, I. & Takahashi, E. Spatiotemporal relationship of brain pathways during human fetal development using high-angular resolution diffusion mr imaging and histology. Front. neuroscience 11, 348 (2017).

33. Aganj, I. et al. Reconstruction of the orientation distribution function in single-and multiple-shell q-ball imaging within constant solid angle. Magn. resonance medicine 64, 554–566 (2010).

34. Rokem, A. et al. Evaluating the accuracy of diffusion mri models in white matter. PloS one 10, e0123272 (2015).

35. Christiaens, D. et al. Scattered slice shard reconstruction for motion correction in multi-shell diffusion mri. Neuroimage 225, 117437 (2021).

36. Jones, D. K. Determining and visualizing uncertainty in estimates of fiber orientation from diffusion tensor mri. Magn. Reson. Medicine: An Off. J. Int. Soc. for Magn. Reson. Medicine 49, 7–12 (2003).

37. Yuan, Y., Zhu, H., Ibrahim, J. G., Lin, W. & Peterson, B. S. A note on the validity of statistical bootstrapping for estimating the uncertainty of tensor parameters in diffusion tensor images. IEEE transactions on medical imaging 27, 1506–1514 (2008).

38. Jeong, H.-K. & Anderson, A. W. Characterizing fiber directional uncertainty in diffusion tensor mri. Magn. Reson. Medicine: An Off. J. Int. Soc. for Magn. Reson. Medicine 60, 1408–1421 (2008).

39. Karimi, D., Warfield, S. K. & Gholipour, A. Calibrated diffusion tensor estimation. arXiv preprint arXiv:2111.10847 (2021).

40. Ulfig, N., Nickel, J. & Bohl, J. Monoclonal antibodies smi 311 and smi 312 as tools to investigate the maturation of nerve cells and axonal patterns in human fetal brain. Cell tissue research 291, 433–443 (1998).

41. Huang, H. et al. White and gray matter development in human fetal, newborn and pediatric brains. Neuroimage 33, 27–38 (2006).

42. Judaš, M. et al. Structural, immunocytochemical, and mr imaging properties of periventricular crossroads of growing cortical pathways in preterm infants. Am. journal neuroradiology 26, 2671–2684 (2005).

43. Kostović, I., Išasegi, I. Ž. & Krsnik, Ž. Sublaminar organization of the human subplate: developmental changes in the distribution of neurons, glia, growing axons and extracellular matrix. J. Anat. 235, 481–506 (2019).

44. Skare, S. et al. Condition number as a measure of noise performance of diffusion tensor data acquisition schemes with mri. J. magnetic resonance 147, 340–352 (2000).

45. Deprez, M. et al. Higher order spherical harmonics reconstruction of fetal diffusion mri with intensity correction. IEEE transactions on medical imaging 39, 1104–1113 (2019).

46. Kebiri, H. et al. Through-plane super-resolution with autoencoders in diffusion magnetic resonance imaging of the developing human brain. Front. Neurol. 13, 765 (2022).

47. Nath, V. et al. Deep learning reveals untapped information for local white-matter fiber reconstruction in diffusion-weighted mri. Magn. resonance imaging 62, 220–227 (2019).

48. Pietsch, M. et al. A framework for multi-component analysis of diffusion mri data over the neonatal period. Neuroimage 186, 321–337 (2019).

49. Bastiani, M. et al. Automated processing pipeline for neonatal diffusion mri in the developing human connectome project. Neuroimage 185, 750–763 (2019).

50. Jha, R. R., Kumar, B. R., Pathak, S. K., Bhavsar, A. & Nigam, A. Trganet: Transforming 3t to 7t dmri using trapezoidal rule and graph based attention modules. Med. Image Analysis 102806 (2023).

51. Yao, T. et al. A unified single-stage learning model for estimating fiber orientation distribution functions on heterogeneous multi-shell diffusion-weighted mri. arXiv preprint arXiv:2303.16376 (2023).

52. Pinto, M. S. et al. Harmonization of brain diffusion mri: Concepts and methods. Front. Neurosci. 14, 396 (2020).

53. Elaldi, A., Dey, N. & Gerig, G. e(3) × so(3)-equivariant networks for spherical deconvolution in diffusion mri. In Medical Imaging with Deep Learning.

54. Fogtmann, M. et al. A unified approach to diffusion direction sensitive slice registration and 3-d dti reconstruction from moving fetal brain anatomy. IEEE transactions on medical imaging 33, 272–289 (2013).

55. Jakab, A. et al. Disrupted developmental organization of the structural connectome in fetuses with corpus callosum agenesis. Neuroimage 111, 277–288 (2015).

56. Marami, B. et al. Temporal slice registration and robust diffusion-tensor reconstruction for improved fetal brain structural connectivity analysis. NeuroImage 156, 475–488 (2017).

57. Jakab, A., Tuura, R., Kellenberger, C. & Scheer, I. In utero diffusion tensor imaging of the fetal brain: a reproducibility study. NeuroImage: Clin. 15, 601–612 (2017).

58. Pietsch, M., Christiaens, D., Hajnal, J. V. & Tournier, J.-D. dstripe: slice artefact correction in diffusion mri via constrained neural network. Med. Image Analysis 74, 102255 (2021).

59. Veraart, J., Fieremans, E. & Novikov, D. S. Diffusion mri noise mapping using random matrix theory. Magn. resonance medicine 76, 1582–1593 (2016).

60. Tustison, N. J., Avants, B. B., Cook, P. A., Zheng, Y. et al. N4itk: improved n3 bias correction. IEEE transactions on medical imaging 29, 1310–1320 (2010).

61. Gholipour, A. et al. A normative spatiotemporal mri atlas of the fetal brain for automatic segmentation and analysis of early brain growth. Sci. reports 7, 476 (2017).

62. Hrabač, P. et al. The zagreb collection of human brains: entering the virtual world. Croat. Med. J. 59, 283 (2018).

63. Ronneberger, O. et al. U-net: Convolutional networks for biomedical image segmentation. In MICCAI, 234–241 (Springer, 2015).

64. Agarap, A. F. Deep learning using rectified linear units (relu). arXiv preprint arXiv:1803.08375 (2018).

65. Srivastava, N., Hinton, G., Krizhevsky, A., Sutskever, I. & Salakhutdinov, R. Dropout: a simple way to prevent neural networks from overfitting. The journal machine learning research 15, 1929–1958 (2014).

66. Garyfallidis, E. et al. Dipy, a library for the analysis of diffusion mri data. Front. neuroinformatics 8 (2014).

67. Kingma, D. P. & Ba, J. Adam: A method for stochastic optimization. arXiv preprint arXiv:1412.6980 (2014).

68. Schilling, K. G. et al. Histological validation of diffusion mri fiber orientation distributions and dispersion. Neuroimage 165, 200–221 (2018).

69. Kunz, N. et al. Assessing white matter microstructure of the newborn with multi-shell diffusion mri and biophysical compartment models. Neuroimage 96, 288–299 (2014).

70. Tournier, J.-D., Calamante, F. & Connelly, A. Mrtrix: diffusion tractography in crossing fiber regions. Int. journal imaging systems technology 22, 53–66 (2012).

71. Chung, S., Lu, Y. & Henry, R. G. Comparison of bootstrap approaches for estimation of uncertainties of dti parameters. NeuroImage 33, 531–541 (2006).

72. Whitcher, B., Tuch, D. S., Wisco, J. J., Sorensen, A. G. & Wang, L. Using the wild bootstrap to quantify uncertainty in diffusion tensor imaging. Hum. brain mapping 29, 346–362 (2008).

73. Zhu, T., Liu, X., Connelly, P. R. & Zhong, J. An optimized wild bootstrap method for evaluation of measurement uncertainties of dti-derived parameters in human brain. Neuroimage 40, 1144–1156 (2008).

